# Conservation of energetic pathways for electroautotrophy in the uncultivated candidate order *Tenderiales*

**DOI:** 10.1101/2022.05.06.490989

**Authors:** Brian J. Eddie, Lina J. Bird, Claus Pelikan, Marc Mussmann, Clara Martinez-Perez, Princess Pinamang, Anthony P. Malanoski, Sarah M. Glaven

## Abstract

Electromicrobiology can be used to understand extracellular electron uptake in previously undescribed chemolithotrophs. Enrichment and characterization of the uncultivated electroautotroph “*Candidatus* Tenderia electrophaga” using electromicrobiology led to the designation of the order *Tenderiales*. Representative *Tenderiales* metagenome assembled genomes (MAGs) have been identified in a number of environmental surveys, yet a comprehensive characterization of conserved genes for extracellular electron uptake has thus far not been conducted. Using comparative genomics we identified conserved orthologous genes within the *Tenderiales* and nearest neighbor orders important for extracellular electron uptake based on a previously proposed pathway from “*Ca*. Tenderia electrophaga”. The *Tenderiales* contained a conserved cluster we designated *uetABCDEFGHIJ*, which encodes proteins containing features that would enable transport of extracellular electrons to cytoplasmic membrane bound energy transducing complexes such as two conserved cytochrome *cbb*_*3*_ oxidases. For example, UetJ is predicted to be an extracellular undecaheme *c*-type cytochrome that forms a heme wire. We also identified clusters of genes predicted to facilitate assembly and maturation of electron transport proteins, as well as cellular attachment to surfaces. Autotrophy among the *Tenderiales* is supported by the presence of carbon fixation and stress response pathways that could allow cellular growth by extracellular electron uptake. Key differences between the *Tenderiales* and other known neutrophilic iron oxidizers were revealed, including very few Cyc2 genes in the *Tenderiales*. Our results reveal a possible conserved pathway for extracellular electron uptake and suggests the *Tenderiales* have an distribution unlimited ecological role coupling metal or mineral redox chemistry and the carbon cycle in marine and brackish sediments.

**Importance:** Electromicrobiology enables enrichment and identification of chemolithotrophic bacteria capable of extracellular electron uptake to drive energy metabolism and CO_2_ fixation. The recently described order *Tenderiales* contains the uncultivated electroautotroph “*Candidatus* Tenderia electrophaga”. The “*Ca*. Tenderia electrophaga” genome contains genes proposed to make up a previously undescribed extracellular electron uptake pathway. Here we use comparative genomics to show that this pathway is well conserved among *Tenderiales* spp. recovered by metagenome assembled genomes. This conservation extends to near neighbors of the *Tenderiales*, but not to other well-studied chemolithotrophs including iron and sulfur oxidizers. Our findings suggest that extracellular electron uptake may be pervasive among the *Tenderiales* and the geographic location from which metagenome assembled genomes were recovered offers clues to their natural ecological niche.

## Introduction

Chemolithotrophic bacteria contribute to key ecological processes, such as metal-cycling and carbon fixation, yet laboratory isolation and cultivation remain elusive hindering our understanding of their distribution and physiology. Electromicrobiology, the study of the movement of electrons into and out of microbial cells using electrochemistry, can be used to enrich and characterize chemolithotrophic bacteria capable of direct extracellular electron uptake. “*Candidatus* Tenderia electrophaga” was enriched on an electrode from seawater and is proposed to couple direct electron uptake with reduction of O_2_ and CO_2_ fixation for growth (1-3). Originally classified as a member of the *Chromatiales* by 16S rRNA gene (16S) sequence homology, “*Ca*. Tenderia electrophaga” was moved to its own order based upon a more detailed protein sequence based phylogeny (4, 5). The *Tenderiales* are now a recognized order within the *Gammaproteobacteria* (4), however, their relatedness to each other, including conservation of electron transfer proteins, and with other chemolithotrophs has not been determined.

Members of the *Tenderiales* have been identified from a number of environments by metagenomic surveys, with several complete or nearly complete metagenome assembled genomes (MAGs) being recovered from marine and estuarine sediments, hydrothermal vents, and floodplain sediments (4, 6, 7). Closely allied 16S sequences have been identified as probable sulfur-oxidizing symbionts of metazoans, although “*Ca*. Tenderia electrophaga” possesses only a limited suite of known sulfur oxidation enzymes (1). A proposed electron uptake pathway from “*Ca*. Tenderia electrophaga” has been described from genomics, transcriptomics, and proteomics data when grown on an electrode (2, 8), but the extent to which it is distributed across other bacterial species is unknown.

Given the emergence of publically available metagenomic data sets that include putative *Tenderiales* spp. we used comparative genomics to further explore the distribution of putative electron transfer proteins from “*Ca*. Tenderia electrophaga”. We first determined the phylogenetic relationships within the *Tenderiales* order and to closely related orders and other bacteria capable of electron uptake. We then identified orthologous genes among these different groups and looked for those that were 1) previously identified as part of the “*Ca*. Tenderia electrophaga” electron uptake pathway, or 2) encoded proteins with potential roles in electron transfer or electrode biofilms. We found several clusters of genes in addition to the previously noted undecaheme cluster that are likely involved in electron uptake, including a novel porin-cytochrome complex and associated cytochromes, orthologs of the cytochrome cbb_3_ oxidase complex, and a conserved hexaheme cytochrome. Comparison of the proposed electron uptake pathways from “*Ca*. Tenderia electrophaga” to more recently identified members of the *Tenderiales* and other chemolithotrophs offers potential biomarkers for their proposed biogeochemical role in oxidizing insoluble extracellular electron donors.

## Results

### Phylogeny of the *Tenderiales*

A selection of 127 genomes and metagenome-assembled genomes (MAGs) (Table S1) were used to establish the phylogenetic relationship among the *Tenderiales*, as well as between the *Tenderiales* and related orders or species suspected of chemolithoautotrophy via extracellular electron uptake. The *Tenderiales* in-group consisted of publicly available MAGs within the *Tenderiales* order defined by the Genome Taxonomy Database (GTDB) (4) and unpublished *Tenderiales* MAGs collected at a field site in Normandy, France {Martinez-Pérez and Mußmann unpublished} (Table 1). Outgroup genomes and MAGs (hereafter often refered to simply as genomes) were selected from the next closest orders in the GTDB taxonomy, from the NCBI *Chromatiales* taxonomy, and from known iron and sulfur-oxidizing bacteria based upon prior observations that several genes found in “*Ca*. Tenderia electrophaga” have been implicated in iron or sulfur oxidation in other organisms (9).

**Table 1.** Genomes and MAGs used in this study

| Bin Id | Name | CheckM completeness | Assembled length | # contigs | Environment | Reference/ Bioproject |
| --- | --- | --- | --- | --- | --- | --- |
| GCA_011330695 | Ca. Tenderia sp. | 70.69 | 2657668 | 659 | Hydrothermal Sediment | (6) |
| FCS_M_Gamma_047 | MAG FCS_M_Gamma_047 | 94.36 | 4775750 | 250 | Normandy magnetite |  |
| FCS_M_Gamma_048.1 | Ca. Tenderia electrophaga | 93.43 | 3442975 | 149 | Normandy magnetite |  |
| 3300032254_18 | Not published metagenome bin | 89.96 | 2901507 | 100 | Damariscotta River Estuary (USA) | Emerson PRJNA539541 |
| FO_M_Gamma_043.1 | MAG FO_M_Gamma_043.1 | 85.59 | 3075132 | 243 | Normandy magnetite |  |
| FCS_M_Gamma_047.1 | MAG FCS_M_Gamma_047.1 | 84.28 | 3032772 | 134 | Normandy magnetite |  |
| 3300037424_47 | Not published metagenome bin | 85.6 | 2906281 | 319 | Floodplain sediments, Riverton, Wyoming, USA | (7) |
| FO_M_Gamma_044 | MAG FO_M_Gamma_044 | 80.47 | 2414050 | 463 | Normandy magnetite |  |
| 3300032260_22 | Not published metagenome bin | 86.71 | 2396328 | 216 | Coastal sediment, Maine (USA) | Emerson PRJNA539527 |
| FO_M_Gamma_046.1 | MAG FO_M_Gamma_046.1 | 74.52 | 2486236 | 246 | Normandy magnetite |  |
| FCS_M_Gamma_048.2 | MAG FCS_M_Gamma_048.2 | 64.99 | 1816566 | 88 | Normandy magnetite |  |
| FO_M_Gamma_045 | MAG FO_M_Gamma_045 | 56.73 | 2313911 | 624 | Normandy magnetite |  |
| 3300032373_40 | Not published metagenome bin | 50 | 2221472 | 280 | Damariscotta River Estuary (USA) | Emerson PRJNA539537 |
| GCA_011375055 | Ca. Tenderia sp. | 37.93 | 1287708 | 460 | Hydrothermal Sediment (Pacific) | (6) |
| CP013099 | Ca. Tenderia electrophaga | 99.63 | 3763866 | 2 | Marine sediment (Atlantic) | (5) |
| GCA_002415485 | Proteobacteria bacterium UBA5122 | 92.33 | 2962318 | 88 | Noosa River Estuary sediment (Australia) | (4) |

“*Ca*. Tenderia electrophaga” NRL1 was the most complete MAG (99.63%) within the *Tenderiales* due to long-read sequencing from a highly enriched community. All other MAGs came from short read metagenomic sequencing projects, and were made up of 88 – 624 contigs with a mean N50 of 25,231. The mean GC content of all short read *Tenderiales* MAGs was 51.66%, with an average genome length of 2.78 Mbp. The mean completeness of the MAGS based upon conserved genes was 77%, which allowed us to extrapolate a mean genome length of 3.6 Mbp, which agrees well with the length of the closed genome of strain NRL1 (3.66 Mbp).

A phylogenetic tree was constructed using an approximately 5000 amino acid alignment of conserved protein domains identified by the tool GTDB-tk (Figure 1). This analysis placed all 16 of the *Tenderiales* into a single well supported clade most closely related to the putative order SZUA-140 identified in the GTDB. This genomic comparison was complemented by a second phylogenetic tree constructed from the 31 nearly full length *Tenderiales* 16S rRNA gene sequences (>1350) in the ARB-Silva non-redundant database and 16S sequences extracted from the genomes and MAGs (Supplemental Figure S1), although it was not possible to determine the relationship between the *Tenderiales* and group 2 MAGs due to the lack of high quality 16S rRNA genes identified in these MAGs.

**Figure 1.**
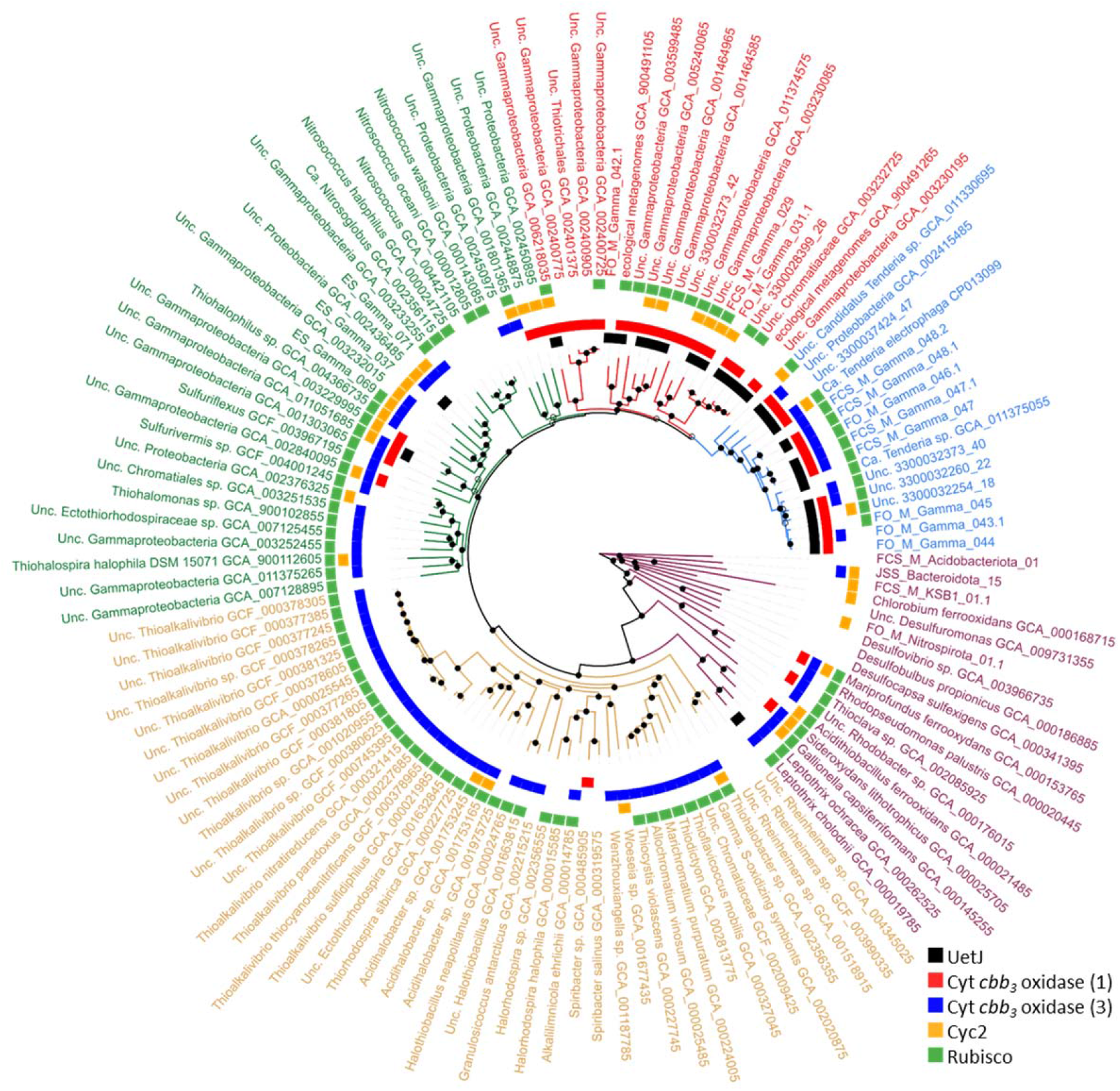
Phylogenetic tree constructed using alignments of conserved proteins collected by CheckM from metagenome bins and outgroup genomes. Group 1 – *Tenderiales*, blue; Group 2 – GTDB neighboring orders, red – 20; Group 3 – GTDB next nearest neighboring orders, green – 30; Group 4 – *Chromatiales*, olive - 43; Group 5 – Functional outgroup organisms, Purple – 18. Taxonomic Order names are from the GTDB. Open circles indicated bootstrap support of >50% for the node, filled circles indicate bootstrap support of > 75%. Colored boxes indicate presence of an ortholog for select marker proteins. For cytochrome *cbb*_*3*_ oxidase, the catalytic subunit CcoN was used for the marker protein and for Rubiscos, presence of either a Type I large subunit was used, or type II protein was used.

### Conserved genes within the Tenderiales MAGs

We assigned groupings to genomes used for phylogenetic comparison based on relatedness to the *Tenderiales* in order to describe the distribution of genes for extracellular electron uptake (Figure 1). Group 1 consisted of the *Tenderiales*. Group 2 consisted of the three neighboring orders in the GTDB. Groups 3 and 4 consisted mostly of unclassified *Gammaproteobacteria* and members of the *Chromatiales*, according to the NCBI taxonomy classification. Group 5 consisted primarily of other iron and sulfur oxidizing species and some additional MAGs recovered from Normandy, France. Orthologous genes, genes descended from a common ancestor via speciation event or horizontal transfer but not gene duplication (10), were identified using ProteinOrtho (Supplemental Table S2)(11). A set of orthologous genes identified across genomes is referred to as an orthogroup. Conservation of orthogroups was quantified based upon relatedness to the *Tenderiales* clade in the protein tree (Figure 1). Orthogroups were named using the representative protein identifier from “*Ca*. Tenderia electrophaga” if present (e.g. ALP54624.1), or the first protein identifier in the arbitrary genome input order if an ortholog was not present in “*Ca*. Tenderia electrophaga” (e.g. ACL71705.1) (Supplemental table S2). A core orthogroup genome was constructed using a 75% cutoff, based on a mean completeness of 77% within the *Tenderiales* MAGs, which resulted in 390 genes unique to the *Tenderiales* (group 1), 508 genes shared between the *Tenderiales* and group 2, and 369 genes common to groups 1-4 (Supplemental Figure S2).

### Conserved proteins for extracellular electron uptake

While direct electron uptake has been noted for a range of microbial species (12), no universal conserved genetic markers for proteins involved in this process have been identified. “*Ca*. Tenderia electrophaga” has been proposed to take up electrons from electrodes via several multiheme cytochromes and other predicted periplasmic, outer membrane, and extracellular proteins (1). Therefore, we looked for genes and gene clusters (orthogroups with a conserved gene order) (Figure 2) for these proteins across all five groups.

**Figure 2.**
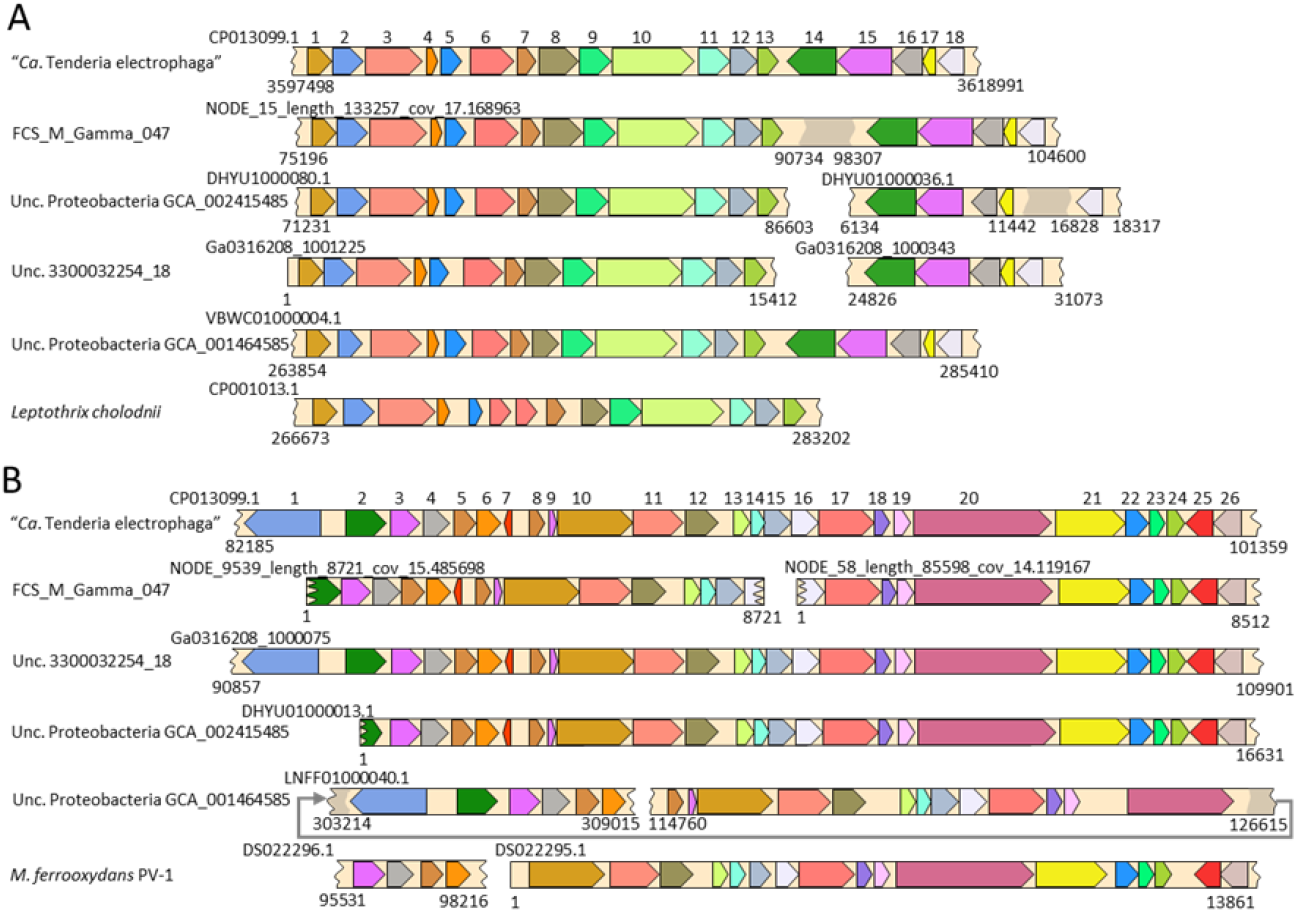
Synteny within conserved genes among the Tenderiales. A) Gene order within the putative EET module found in the *Tenderiales* is highly conserved among a diverse selection of the *Tenderiales* and group 2 organisms, with an outlier found in *Leptothrix cholodnii*. Orthologous genes share the same color, with labels shown for *Ca*. Tenderia electrophaga proteins encoded. 1 – ALP54622.1 Tetraheme cytochrome UetA; 2 - ALP54623.1 peptidylprolyl isomerase UetB; 3 - ALP54624.1 – β-barrel outer membrane protein UetC; 4- ALP54625.1 triheme cytochrome *c*(7) UetD; 5 - ALP54626.1 triheme cytochrome *c*(7) triheme UetE; 6- ALP54627.1 Cupredoxin UetF; 7 - ALP54628.1 triheme cytochrome *c*(7) UetG; 8- ALP54629.1 NHL repeat unit UetH; 9- ALP54908.1 NHL repeat unit <u>U</u>etI; 10 - fig|6666666.657277.peg.3443 Undecaheme cytochrome UetJ; 11- ALP54630.1 *c*-type cytochrome biogenesis protein CcsA; 12 - ALP54631.1 *c*-type cytochrome biogenesis protein CcsB; 13 - ALP54632.1 TPR repeat NrfG; 14 - ALP54633.1 BatD domain protein; 15 - ALP54634.1 BatB von Willebrand family protein; 16 - ALP54635.1 BatA von Willebrand family protein; 17 - ALP54636.1 hypothetical protein; 18 – ALP54637.1 YeaD2 superfamily. B) The region containing cytochrome *cbb*_3_ oxidase 1 and 2 is conserved among the *Tenderiales*, and some more distantly related organisms. 1 - ALP51721.1; 2 - ALP51722.1 di-heme cytochrome; 3 - ALP51723.1 monoheme cytochrome; 4 – ALP51724.1 truncated CcoN cytochrome c oxidase; 5 - ALP51725.1 Thioredoxin; 6 - ALP51726.1; 7 - ALP51727.1; 8 - ALP51728.1; 9 - 657277.peg.91; 10 - ALP51729.1 cytochrome C oxidase CcoN; 11 - ALP51730.1 cytochrome oxidase CcoO; 12 - ALP51731.1; 13 - ALP51732.1; 14 - ALP51733.1; 15 - ALP51734.1 monoheme cytochrome; 16 - ALP51735.1; 17 - ALP51736.1 serine threonine protein kinase; 18 - ALP51737.1 STMD; 19 - ALP51738.1; 20 – ALP54674.1; 21 - ALP51739.1 CcoG *cbb*_*3*_ maturation protein; 22 - ALP51740.1; 23 - ALP51741.1 monoheme cytochrome; 24 - ALP51742.1; 25 - ALP51743.1; 26 - ALP51744.1 truncated CcoN cytochrome oxidase

#### Undecaheme cytochrome, tri-heme cytochrome domains, NHL repeats

We previously noted that genes for an undecaheme and three tri-heme *c*-type cytochromes were collocated on the “*Ca*. Tenderia electrophaga” genome and were differentially expressed during EET depending on the electrode potential (2). These genes were part of a block of genes encoding 18 orthogroups (Supplemental table S4) that were highly conserved among the *Tenderiales* and Group 2 organisms (Figure 1, black squares). Furthermore, the order of genes was remarkably well conserved (Figure 2A), indicating selective pressure for collocation. Here, we name the first ten of these genes *uetABCDEFGHIJ*, for <u>u</u>ndecaheme <u>e</u>lectron <u>t</u>ransfer, and we refer to this region as the *uet* cluster, and include the downstream conserved genes, *ccsAB, nrfG*, and *batABD* as a conserved part of this module (Figure 3).

**Figure 3.**
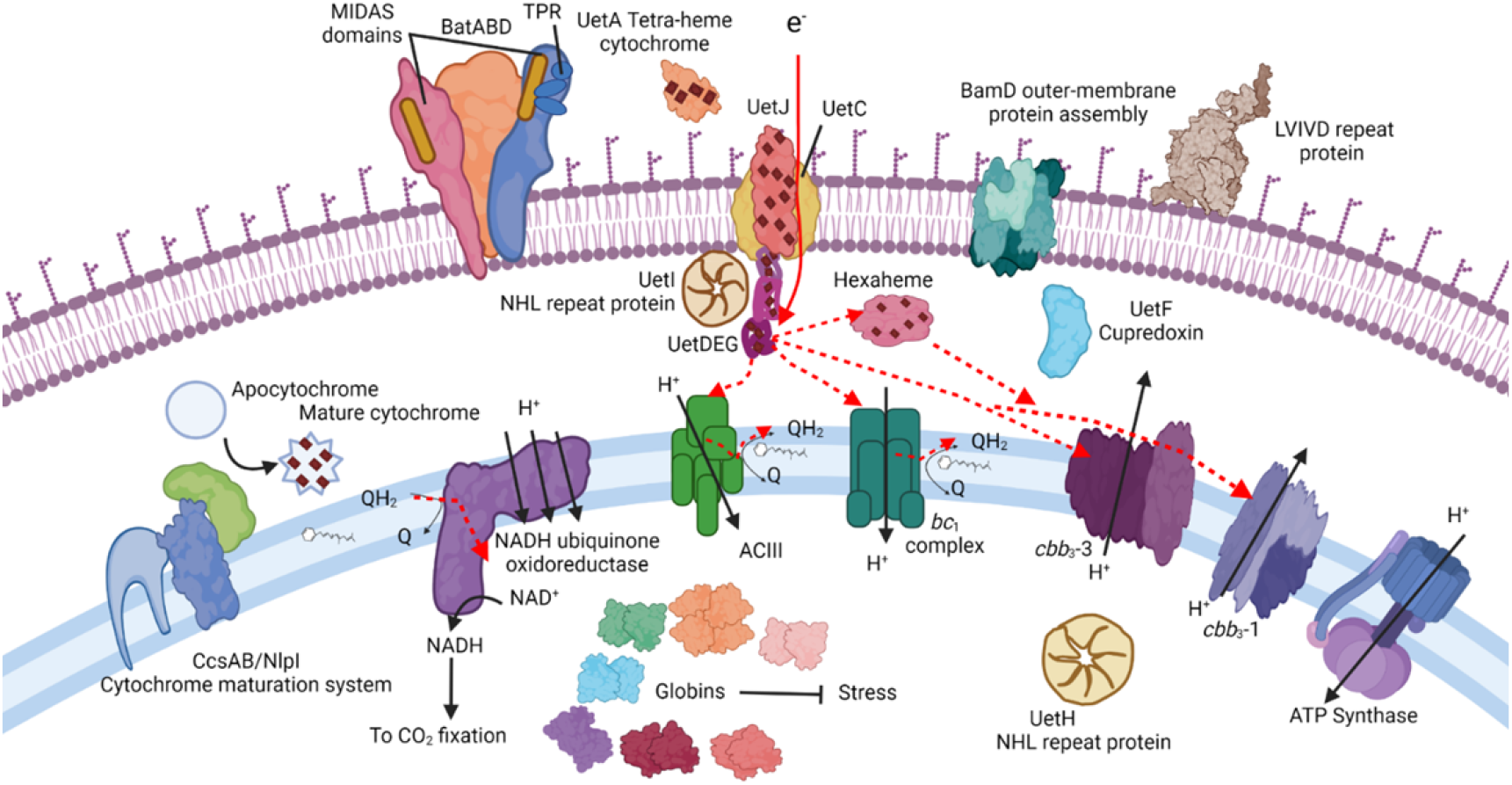
Predicted energetic pathway and auxiliary pathways found in the *Tenderiales*. Predicted electron flow in this model is shown with red arrows. Globins fulfill a stress response function. CcsAB/NlpI are proposed to be required for maturation of Uet cytochromes. The two BatABD complexes and the LVIVD repeat protein likely facilitate attachment to substrates. BamD and associated proteins are predicted to be required for maturation of outer-membrane β-barrel proteins. Presence of individual pathway components in each MAG can be found in supplemental table S2.

The undecaheme protein UetJ (fig|6666666.657277.peg.3443), contained eleven CXXCH heme binding motifs but no other conserved functional domains. It is highly expressed (2), suggesting its designation as a pseudogene by NCBI is an annotation error. *In silico* structural predictions indicated a long, bar shape with the heme binding domains lined up down the center of the protein less than 1 nm apart, which would allow for efficient electron transfer (13) (Supplemental Figure S3). It was predicted to contain a signal peptide for export from the cytoplasm and was predicted to be an extracellular protein by pSort3.0 (14). UetC (ALP54624.0) was a predicted β-barrel protein that could form a pore into which an electron transfer protein is inserted, much as MtrA fits inside MtrB (15). In contrast to MtrB, which contains 26 transmembrane strands, UetC was predicted by PRED-TMBB2 to contain 28 transmembrane strands, which likely gives it a larger pore size (16). This was borne out in the structural model, with a pore of approximately 5 nm diameter (Supplemental Figure S3).

UetDEG (ALP54625.1, ALP54626.1, and ALP54628.1) each contained a *c*(7)-type tri-heme domain previously identified in periplasmic electron transfer proteins from *Geobacter sulfurreducens* (17). The best structurally characterized of the *G. sulfurreducens* proteins, the periplasmic dodecaheme *c*-type cytochrome GSU1996, contains four such tri-heme domains (A, B, C, and D) (17), of which, all three “*Ca*. Tenderia electrophaga” orthologs best match domain C. UetF (ALP54627.1) contains a predicted cupredoxin domain, which have been associated with electron transfer via a Cu^+1^ ion. Proteins with a single cupredoxin domain are often found as soluble electron carriers such as azurin, rusticyanin, and plastocyanin. However UetF was much larger than these proteins and may have a different role here.

UetH and UetI (ALP54629.1 and ALP54908.1) contained NHL domains, a β-propeller structure that is sometimes associated with protein:protein interactions (18). UetH was predicted by PSORTb to be cytoplasmic, with a seven-bladed β-propeller structure, while UetI was predicted to be non-cytoplasmic, with a six-bladed β-propeller structure. An NHL repeat domain containing protein (Gmet_0556) was identified as being important in extracellular iron reduction in *G. metallireducens*, most likely due to cellular adhesion (19) and stabilization of outer membrane cytochrome complexes in *Desulfovibrio ferrophilus* IS5 (20).

#### Cytochrome c maturation and assembly

The predicted orthogroups represented by ALP54630.1 and ALP54631.1 were similar to the type-II cytochrome maturation system proteins CcsA and CcsB, not commonly found in *Gammaproteobacteria*. The orthogroup represented by ALP54632.1 contained tetratricopeptide repeat (TPR) domains and a partial NrfG domain, which have been shown to facilitate the maturation of multiheme cytochromes (21). The presence of this system in addition to the type-I cytochrome maturation system might indicate some unusual aspect of the *Tenderiales*. For example, it has been speculated that Anammox bacteria use the type-II system as a second maturation complex dedicated toward maturation of multiheme cytochromes within the Annamoxosome organelle (22). There is also recent evidence for horizontal gene transfer of cytochrome maturation systems along with EET modules (23).

#### Adhesion and metal-binding domains

The *uet* cluster was adjacent to conserved orthogroups that were homologs of the *Bacteroides* AeroTolerance (BatI) operon (Figure 2A), which can play a role in adhesion. ALP54633.1 contained a BatD domain with a single transmembrane domain, allowing it to be anchored in the outer membrane. In the *batI* operon, *batD* was preceded by a gene for the von Willebrand adhesion factor domain and TPR protein:protein interaction domain containing lipoprotein BatB (ALP54634.1) and BatA von Willebrand factor domain protein (ALP54635.1) (24). The metal ion dependent adhesion site (MIDAS) motif DXSXS was conserved in both proteins near the N-terminal. Originally associated with biofilm formation and adhesion in the presence of Mg^2+^, this domain is associated with anodic EET in the Gram-positive bacterium *Enterococcus faecalis* by moderating some interaction with extracellular iron (25). The *Tenderiales* also contained a second BatABD operon represented by ALP52718.1 – ALP52720.1 that was similar to the one discussed above.

#### Secondary c-type cytochrome containing cluster

The *batI* operon in the “*Ca*. Tenderiales electrophaga” genome was followed by a ten gene type-I cytochrome maturation pathway found in most of the genomes examined here, and a cluster of 9 orthogroups that were conserved in the *Tenderiales* and Group 2 organisms, with only a few representatives found in groups 3-5. Three of these orthogroups may be involved in regulation, with conserved domains for periplasmic ligand-binding sensor domains (ALP54649.1 and ALP54650.1) and a heme containing cyclic di-GMP phosphodiesterase (ALP54651.1). This cluster of genes encodes several potential electron transfer proteins, including a thioredoxin (ALP54648.1) and four multiheme cytochromes. These cytochromes consist of one with a partial CcoP domain (ALP54909.1), a pentaheme (ALP54653.1) and a two tri-heme cytochromes (ALP54654.1 and ALP54655.1). ALP54655.1 also contained a CccA domain, which is implicated in iron oxidation. There was also an NHL domain protein (ALP54653.1), which may play a role in structuring these proteins into a complex.

### Other potential electron transfer proteins

#### Hexaheme cytochrome c

A small cluster of three genes encoding proteins with *c*-type cytochrome heme-binding motifs was well conserved among the *Tenderiales*. Monoheme (ALP52411.1), hexaheme (represented by ALP52412.1) and di-heme cytochromes (ALP52413.1) were identified in 14, 15, and 16 of the 16 *Tenderiales* genomes respectively, but were only collocated with each other in 10 of the 16 genomes. The hexaheme protein was predicted to be localized to the periplasm. Despite the presence of six heme-binding domains, it does not appear to be closely related to other characterized hexaheme cytochromes, such as OmcS from *Geobacter sulfurreducens* or TherJR_1122 from *Therminicola potens* (26, 27). The di-heme cytochrome contains a signal peptide, and was predicted to be extracytoplasmic by pSort3.0, but no further functional prediction was possible. The monoheme cytochrome was predicted to be localized to the cytoplasmic membrane, with four predicted transmembrane helices predicted by Phobius, and a CXXCH heme binding motif that was predicted to be exposed to the periplasm.

#### Cyc2

Cyc2 is a biochemically verified iron oxidase, and is widespread throughout several iron oxidizer lineages (28). “*Ca*. Tenderia electrophaga” was proposed to be an iron-oxidizing bacterium based, in part, on the presence of a Cyc2 (ALP52279.1) (1). This protein was found in an orthogroup of only seven genes. Only one other member of the *Tenderiales* had a member of this orthogroup. Manual Blast searches revealed five more orthogroups with a total of 36 additional proteins, bringing the total to 43 proteins spread across 36 genomes (Figure 1), however, only four *Tenderiales* total had a Cyc2. Because so few Cyc2 homologs were identified within the *Tenderiales* and closely related MAGs (Figure 1), it is unlikely that it plays anything more than an auxiliary role in EET in the *Tenderiales*.

### Electron transport chain

#### Electron transfer to the quinone pool: conservation of the ACIII and bc_1_ complex

In our model, upon entry into the cell through UetJ and one of several electron transfer proteins in the periplasm, electrons must travel to the terminal electron acceptor, O_2_, or via the intramembrane quinone pool to reduce NAD^+^ for CO_2_ reduction (Figure 3). The *Tenderiales* possessed two conserved entry points into the quinone pool for electrons from EET, the alternative complex III (ACIII) and cytochrome *bc*_*1*_ complex. The individual genes for both complexes were present in 9-15 of the genomes, depending on the gene.

The ACIII was originally described in bacteria due to the absence of the *bc*_*1*_ complex (9), so the presence of a full *bc*_*1*_ complex in addition to the ACIII suggests that they may fill separate functional roles, such as forward and reverse electron transport. Such a configuration has been hypothesized to be important to extracellular electron transfer and iron oxidation (2, 29, 30). We found that although the gene order was not always conserved, the gene cluster representing the ACIII complex was found in 11 of the *Tenderiales*, and most of the group 2 organisms and the microaerophilic iron oxidizers.

The genes for the cyt-*bc*_*1*_ complex were also present in most of the *Tenderiales* (13-15 genomes), and the proteins for the Rieske Fe-S subunit and cytochrome *b* were identified as orthologs of proteins in the majority of the outgroup genomes. However, the cytochrome *c*_*1*_ orthologs were split into two different orthogroups, with the *Tenderiales* and about half of the group 2 organisms possessing members of one orthogroup which was uncommon outside of the *Tenderiales*. This protein was similar enough to characterized cytochrome *bc*_*1*_ complex subunits that the Conserved Domain Database identified it as a member of COG2857 ubiquinol-cytochrome *c* reductase, however, a second larger orthogroup of cytochrome *c*_*1*_ proteins was identified in members of groups 3, 4 and 5. This second orthogroup did not contain any representatives from group 1 or 2 genomes, and members of the two orthogroups were mutually exclusive within a genome.

Both the NADH ubiquinone:oxidoreductase (NUOR) and succinate dehydrogenase (SDH) encoding regions were well conserved across all groups. NUOR is likely to be involved in reverse electron transport (2), accepting electrons from quinones and using proton motive force to reduce NAD^+^ to NADH that can be used for anabolic processes within the cell as has been previously reported for a wide range of chemolithoautotrophs, including iron-oxidizing bacteria (29, 31). Indeed, 12 of the 16 *Tenderiales* MAGs, and most of the other genomes as well, encoded at least one Form I Rubisco and the necessary anaplerotic enzymes for CO_2_ fixation via the NADH consuming Calvin-Benson-Bassham cycle (Supplementary Table S4). SDH would allow heterotrophic growth based upon intracellular storage products, such as glycogen.

### Terminal oxidases

We previously reported that “*Ca*. Tenderia electrophaga” encodes three cytochrome *cbb*_*3*_ oxidase complexes, which are typically associated with microaerophilic respiration (2), and these complexes were conserved across the *Tenderiales*. The complex that was orthologous to the best studied representatives of *cbb*_*3*_ oxidase consists of a full *cbb*_*3*_ type cytochrome *c* oxidase (CcoNOQP; ALP52999.1-ALP53002.1). We predicted this to be the main terminal oxidase in the electron transport chain of “*Ca*. Tenderia electrophaga” on the basis of metatranscriptomics (2). It was present in most of the *Tenderiales* MAGs and was also common in other organisms. This region also encoded two conserved small single transmembrane domain (STMD) proteins that were rare outside of the *Tenderiales* (fig|6666666.657277.peg.1551 and fig|6666666.657277.peg.1556). STMDs are small proteins (∼35AA) that have a single transmembrane domain with 5-10 exposed residues at either end. They are involved in assembly of respiratory complexes, but have mostly been studied in a mitochondrial context (32). Genes encoding other *cbb*_*3*_ accessory proteins were conserved within the *Tenderiales* as well, including FixHIS and the cytochrome *c* biogenesis protein DsbD. A gene encoding the ferredoxin FixG was only present in “*Ca*. Tenderia electrophaga” and two other members of the *Tenderiales*, but was relatively common in groups 3, 4, and 5. This arrangement was also common in the endosymbiotic nitrogen fixing bacteria, including *Ensifer meliloti*, which leads to an interesting possibility of horizontal gene transfer that is beyond the scope of this study (33).

A large region of 30 genes containing the previously identified *cbb*_*3*_-1 and *cbb*_*3*_-2 complexes in “*Ca*. Tenderia electrophaga” contained orthologs of 26 proteins with a conserved arrangement (Figure 2B) that were identified in most group 1 and 2 genomes, and were nearly absent in groups 3 and 4. This region was previously identified as conserved in some freshwater iron-oxidizing bacteria (34). Most predicted proteins in this region contain transmembrane helices or are membrane associated, suggesting that it represents a membrane bound complex, with several proteins potentially facilitating assembly of the complex. The two partial *cbb*_*3*_ type cytochrome-*c* oxidases indicated this putative complex is likely involved in the electron transport chain and may generate proton motive force, as a sort of alternative complex IV. Conservation of these genes within the *Tenderiales* and group 2 organisms suggests their function is also conserved. Their distribution was similar to the *uetA-J* genes, indicating a functional link between these two modules, but it is also possible this was an artifact of relatedness between the genomes, because *cbb*_*3*_-1 and *cbb*_*3*_-3 were slightly more broadly distributed.

### Other conserved regions of note

#### Conserved Globin Proteins Indicate Stress Response

We noted several conserved proteins related to stess response and biofilm formation, which are crucial processes for EET on electrodes. Seven globin proteins (39), here named globin-1 through globin-7, formed conserved orthogroups in *Tenderiales*, but were rare in other groups (Table S4). Globin-1 (ALP51974.1) was found almost exclusively in *Tenderiales* and was adjacent to a 2-oxoglutarate-Fe(II) oxygenase (ALP51975.1), while globin-2 (ALP52420.1) is found in most *Tenderiales* and about half of their nearest neighbor orders. Some globins were also found in known iron-oxidizing bacteria, including the triple globin domain protein globin-3 (Ga0316208_100021114) - *Sideroxydans lithotrophicus* ES-1, globin-6 (ALP54872.1) - *Mariprofundus ferrooxidans* PV1, and globin-7 (Ga0316208_100045921) - *Rhodopseudomonas palustris*, and *Acidothiobacillus ferrooxidans*. Globin-5 (ALP54375.1) was only conserved at 68.75%, but it was significantly more highly expressed under the more stressful condition in a biocathode metatranscriptome (2).

#### Uncharacterized surface and attachment proteins

The *Tenderiales* contained a number of unique, uncharacterized surface associated proteins that may be important for biofilm formation on solid electron donors. These were identified by searching for domains involved in adhesion and screened for abundance (≥75%) in the *Tenderiales* and rarity (<50%) in groups 3 and 4 (Supplemental table S3). Five orthogroups were found which contained adhesion related histidine kinase or diguanylate-cyclase domains. These could regulate attachment as found in other electroactive biofilms (35, 36).

Several other orthogroups contained domains that suggested a role as extracellular structural components or secretion of exopolysaccharides. Thirteen *Tenderiales* had orthogroup ALP52417.1, containing a bacterial polysaccharide deacetylase PgaB domain, required for export of biofilm forming polysaccharides in *Staphylococcus epidermidis* (37). In five of the *Tenderiales*, this was paired with an orthogroup containing a PulE pilin secretion domain which was not found in any other groups (ALP52418.1). The orthogroup represented by ALP54212.1 was found only in the *Tenderiales*, and contains an LVIVD repeat domain, which is associated with the cell surface of bacteria and archaea (38). This protein may form a β-propeller structure, which can be associated with protein:protein interactions (38). It was adjacent to the orthogroup ALP54213.1, which was also conserved in the *Tenderiales*, with no functional predictions. However, it does have a predicted signal peptide for export from the cytoplasm.

A cluster of four orthogroups that may export and assemble extracellular proteins was mostly confined to the *Tenderiales*. Orthogroup ALP54158.1 had a BamD domain associated with folding and insertion of outer membrane β-barrel proteins (39), including the five TPR repeats required for protein function. This orthogroup was well conserved in the *Tenderiales*, and less well conserved in the other groups here, with representation in 35% of group 2 organisms, and 27% of group 3. It was followed by ALP54159.1, which contains three TPR repeats and was only found in the *Tenderiales* and two group 2 genomes, and ALP54160.1 which was only found in the *Tenderiales*, and did not have any conserved domains (40). The orthogroup represented by ALP54162.1 was similar to the β-barrel protein BamA which is necessary for exporting the protein to be folded. However, this was only present in half of the *Tenderiales* genomes.

## Discussion

Here, comparative genomics was used to understand the metabolic potential of the *Tenderiales*. Based upon the conservation of genes for Rubisco and identification of genes that may be involved in EET, members of this order are likely all chemolithoautotrophs, using extracellular electron donors to fix CO_2_. We described a key gene cluster, the *uet* cluster, conserved throughout the *Tenderiales* and several closely related orders, that encodes an extracellular undecaheme *c*-type cytochrome along with a number of other genes that likely play a role in EET and biofilm formation and maintenance. This *uet* cluster has not been implicated in EET in other bacterial lineages, but encoded proteins appear to follow the same structural rules identified in EET complexes, such as MtrCAB and PioAB, namely an outer membrane β-barrel protein to provide a conduit into which multiheme cytochromes are inserted to transfer electrons across the outer membrane (41). While it is not possible to determine the precise role of each protein from bioinformatics methods alone, protein localization prediction tools suggest an extracellular location for the tetraheme cytochrome UetA and the undecaheme cytochrome UetJ, and a periplasmic localization for the triheme cytochromes UetD, UetE, and UetG. All five of these proteins have predicted structures that place the hemes in a linear arrangement, ideal for transferring electrons over long distances. The predicted extracellular cytochromes UetJ and UetA may interact, with one acting as a transmembrane electron conduit. It is also possible that one or more of them forms long conductive polymers like those in *G. sulfurreducens* (27). This might enable electron transfer in the near-cell environment, and may be responsible for the remarkable conductivity of a biocathode community containing “*Ca*. Tenderia electrophaga” (42).

Electrons entering the cell through such a pathway are then are shuttled across the periplasm to the cytoplasmic membrane, where several options for energy conservation await. The *Tenderiales* contain several conserved modules that give them the ability to grow in a microaerophilic environment, such as two *cbb*_*3*_-type terminal oxidases (43), indicating metabolic flexibility as seen in other organisms (44-46). Based upon the previous metatranscriptomic results, the main terminal oxidase in the reduction of O_2_ is most likely *cbb*_*3*_ oxidase-3, but its role relative to the other terminal oxidase is unknown. The *Tenderiales* posess several electron transport chain components that are also found in iron-oxidizing bacteria, such as ACIII (47) and a cytochrome *bc*_*1*_ complex with a unique cyt-*c*_*1*_ protein that may indicate that it interacts with a unique redox partner, possibly for reverse electron transport. The presence of seven conserved globin genes in the *Tenderiales* may be a response to various oxidative or nitrosative stresses experienced by these organisms, similar to their proposed role in *M. ferrooxydans* (34). The psychrophile *Pseudoalteromonas haloplanktis* contains four constitutively expressed globins that have been identified as playing distinct roles in reactive oxygen and nitrogen stress response (48, 49). The presence of a 2-oxoglutarate Fe(II) oxygenase is intriguing, because α-keto-acids are known to have a peroxynitrite detoxifying effect, suggesting a connection between this globin orthogroup and nitrosative stress (50).

Biofilm formation is a key component of survival on an insoluble extracellular electron donor, and we see evidence for several unique orthogroups that may be involved in formation of biofilms. These range from BatABD proteins, which are similar to proteins that have been identified as critical for EET in *Streptococcus* (25), to polysaccharide secretion, which is necessary for forming the matrix of a biofilm. We also see a conserved orthogroup that is likely involved in assembly of outer-membrane complexes, and an LVIVD-domain protein which may be involved in stabilizing interactions with extracellular proteins.

Microbial community surveys using the 16S rRNA gene and higher resolution surveys using metagenomics have revealed the astonishing diversity of the bacterial world, and allowed us to see that there is a considerable amount of diversity within the *Tenderiales*, including at least two major clades consisting of three genera. Several of the representative sequences were obtained from communities present on potential donors for EET, such as iron sulfide minerals (51) supporting the prediction that they are capable of extracellular electron uptake. Most *Tenderiales* MAGs are from marine or brackish environments, but two were from river sediments(7), along with some of the *Tenderiales* 16S sequences (52). The most closely related MAGs were obtained from environments including Normandy magnetite and hydrothermal vents, which might indicate a potential for an electroautotrophic lifestyle for these additional orders as well.

Furthermore, the high level of conservation of some predicted EET complexes and proteins within the *Tenderiales*, and some of the most closely related orders suggests an important role for these components within cellular metabolism. We are not aware of any predicted metabolism for any members of the group 2 orders, but the presence of the *uetABCDEFGHIJ* region in these MAGS suggests a shared function that we predict to be oxidation of extracellular electron donors. The closest cultivated strains to the *Tenderiales*, such as the *Nitrosococcales* and *Thiohalomonadales*, are chemolithoautotrophs, and the uncultivated organisms found in groups 1 and 2 also appear to be capable of chemolithoautotrophic growth, with Rubisco genes and potential EET conduits. All this supports an ecological role for the *Tenderiales* in oxidizing insoluble electron donors such as iron minerals and fixing carbon within sediment.

The genes we have identified here expand our toolkit for beginning the process of engineering organisms that derive a biosynthetic advantage from using a bioelectrochemical system for redox balancing or supplemental energy to increase production efficiency (53-55). We have identified conserved protein coding genes in the uncultivated order *Tenderiales*, including some which we propose form a novel conduit for EET. Control of the processes involved in formation and maintenance of biofilms, as well as mitigation of stresses involved in this lifestyle will be key to further development of these organisms for biotechnology.

## Methods

### Selection and curation of genomes for analysis

Genome sequences, predicted protein sequence, and gene feature files for in-group and out-group genomes were downloaded in May of 2020 from NCBI (https://www.ncbi.nlm.nih.gov) if available. Genomes not available on NCBI were downloaded from the Integrated Microbial Genome database and updated in April of 2021 via release 202 of GTDB (56). Unpublished MAGs were annotated using RAST to generate the required files (57).

The NCBI and RAST annotation pipelines mostly identified the same proteins, but to better standardardize annotations between methods, while retaining relevant NCBI identifiers for previously published genomes, all genomes were reannotated using the RAST pipeline and the two annotations were merged. For proteins predicted by both methods, the original was retained. If reannotation predicted a protein where the original annotation predicted a pseudogene or no gene, the RAST annotation took precedence. Proteins identified by RAST have a gene identifier beginning with “fig”.

CheckM (58) was used to check genomes for completeness, contamination, and other general genome statistics (Table S1). The tool GTDB-tk was used to identify genomes according to the GTDB taxonomy based upon conserved genes (Table S1) (59). Small subunit ribosomal RNAs (SSU rRNAs) were identified within genomes and MAGs using the tool Ssu_finder. Additional non-redundant, near full-length (>1350 bp) SSU-rRNA sequences for the *Tenderiales* were obtained from the Silva nr database (60). This yielded 27 additional sequences.

### Phylogenetic analysis of the *Tenderiales*

A protein based phylogenetic reconstruction was created with conserved single copy protein alignments obtained using CheckM. IQ-Tree (61) was used to reconstruct the phylogenetic tree using a maximum likelihood algorithm with the Le and Gascuel model with frequencies and 10 rate categories (62). Confidence in branching order was assessed using 1000 ultrafast approximate bootstraps. Phylogenetic trees were visualized using Iroki tree viewer (63). The protein based phylogenetic tree was used to separate genomes into groups for further analysis.

### Identification of Conserved Orthologous Proteins

Orthologous predicted proteins were identified using ProteinOrtho v6.0.30 with the default settings and the PoFF extension. The default settings were highly conservative and resulted in splitting some clusters of proteins. Orthologous proteins that have been previously identified as being part of larger groups with different subtypes, i.e. Rubisco proteins and Cyc2 proteins, were identified by using Blast to connect some of the initial clusters of orthologs that were more distantly related for final groupings reported in the paper (11). Conservation of orthogroups (10) was quantified for groups 1-5 defined above. We then identified the *Tenderiales* core genome consisting of genes for predicted proteins that are conserved in at least 75% of *Tenderiales* MAGs.

Synteny between genomes was visualized using SimpleSynteny (64) to identify protein encoding genes that are conserved in order between genomes. The four *Tenderiales* MAGs selected for this were chosen based upon completeness and taxonomic coverage. For the putative electron transfer complex proteins, a MAG from a drinking water system metagenome(65) and the closed genome of the pure culture *L. cholodnii* (66) were chosen because they are more distantly related taxonomically, but contained the genes associated with this complex.

Additional publicly available bioinformatics tools were used for further characterization of conserved proteins (Supplemental table S4). Conserved domains were identified using the Conserved Domain Database hosted by the NCBI (67). Transmembrane helices were identified using Phobius (68). Outer membrane β-barrel predictions were performed using Pred-TMBB2 (16). Heme binding motifs were identified using the regular expression C.{2,4}CH in Notepad++ (https://notepad-plus-plus.org/). Homologous proteins that have been characterized in some way were identified using PaperBLAST (69).

## Data availability

Annotations used in this study are available on FigShare at doi: 10.6084/m9.figshare.19632867

## Acknowledgements

This work was funded through NRL’s Base 6.1 program and the Office of Naval Research (ONR), N0001422WX00222 to SG and the the Austrian Science Fund (FWF), project ID P31010, awarded to MM.

## Supplemental figures and tables

**Supplemental Figure 1.**
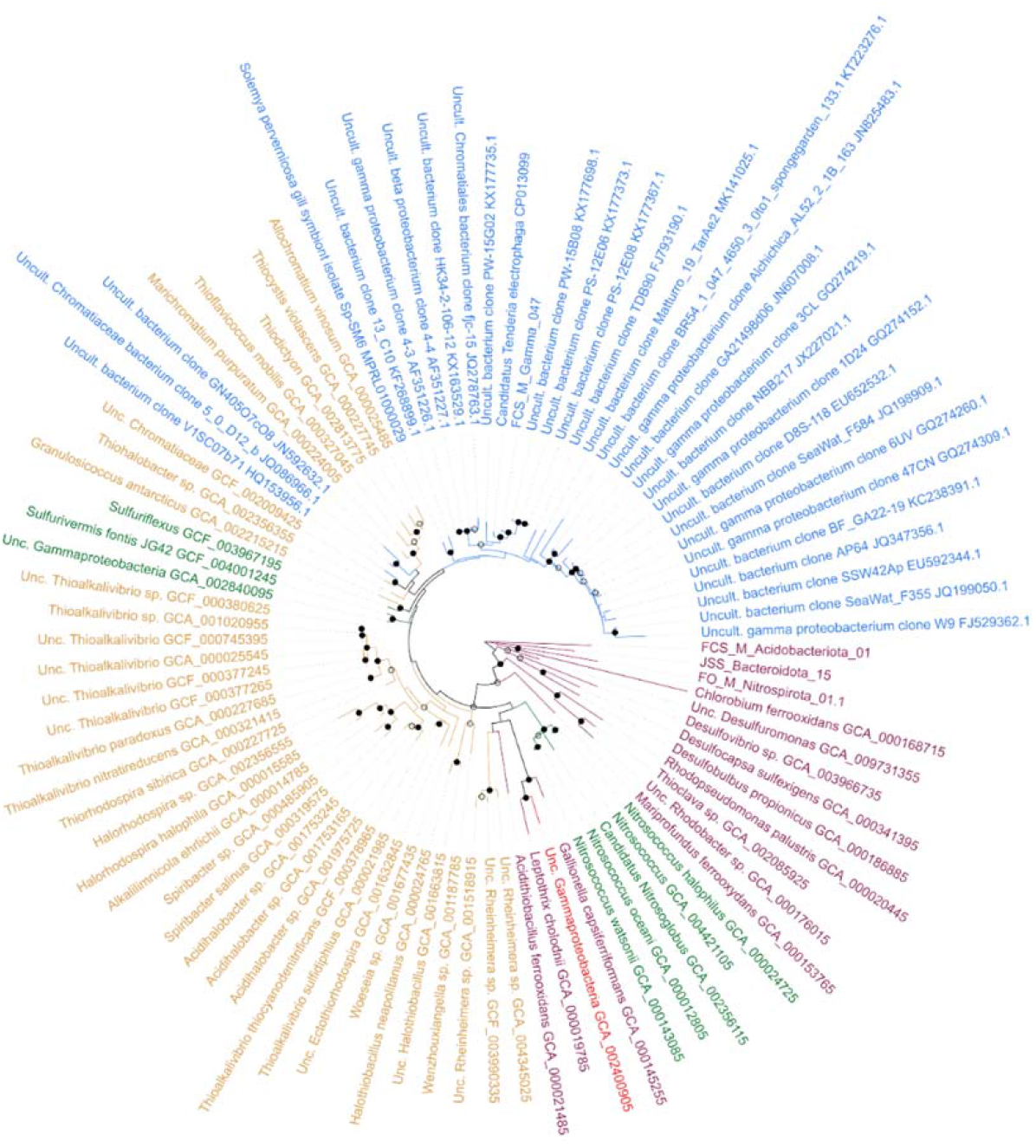
Phylogenetic tree constructed using 16S sequences from the SILVA database and from genomes and MAGs used in this study. Using the Arb-SILVA browser we selected *Tenderiales* sequences from the Silva non-redundant reference database (Silva Ref nr), then removed any that were less than 1375 bp long or had pintail, alignment or sequence quality scores <95% using the Arb-SILVA alignment, classification, tree tool(60). The 31 remaining *Tenderiales* 16S rRNA genes and full length 16S sequences identified in the metagenome assembled genome bins and outgroup genomes listed above were aligned using the SINA alignment tool (70), and trimmed to ends present in >90% of sequences. A phylogenetic tree was inferred using the maximum likelihood algorithm in MEGA version X (71) with the General Time Reversible model for amino acid substitution with 5 gamma categories, based upon results of the MEGA model selection tool. Branching confidence was assessed by performing 500 bootstraps, indicated by an open circle at nodes with >50% support, and filled circles at nodes with >75% support. Due to the difficulty of assembling 16S sequences from the short reads used in most metagenomic studies, many of the assembled MAGs did not contain full length 16S rRNA genes. Scale bar represents 0.1 substitutions per base.

**Supplemental figure S2.**
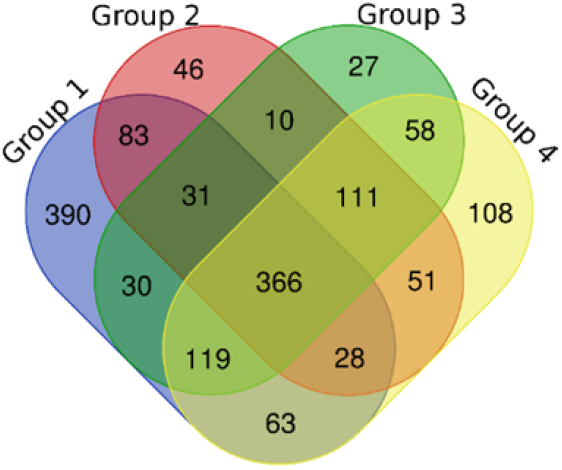
Venn diagram of orthogroups shared among the organism groups. An orthogroup was considered present in an organism group if it was found in at least 75% of representatives of that group.

**Supplemental figure S3.**
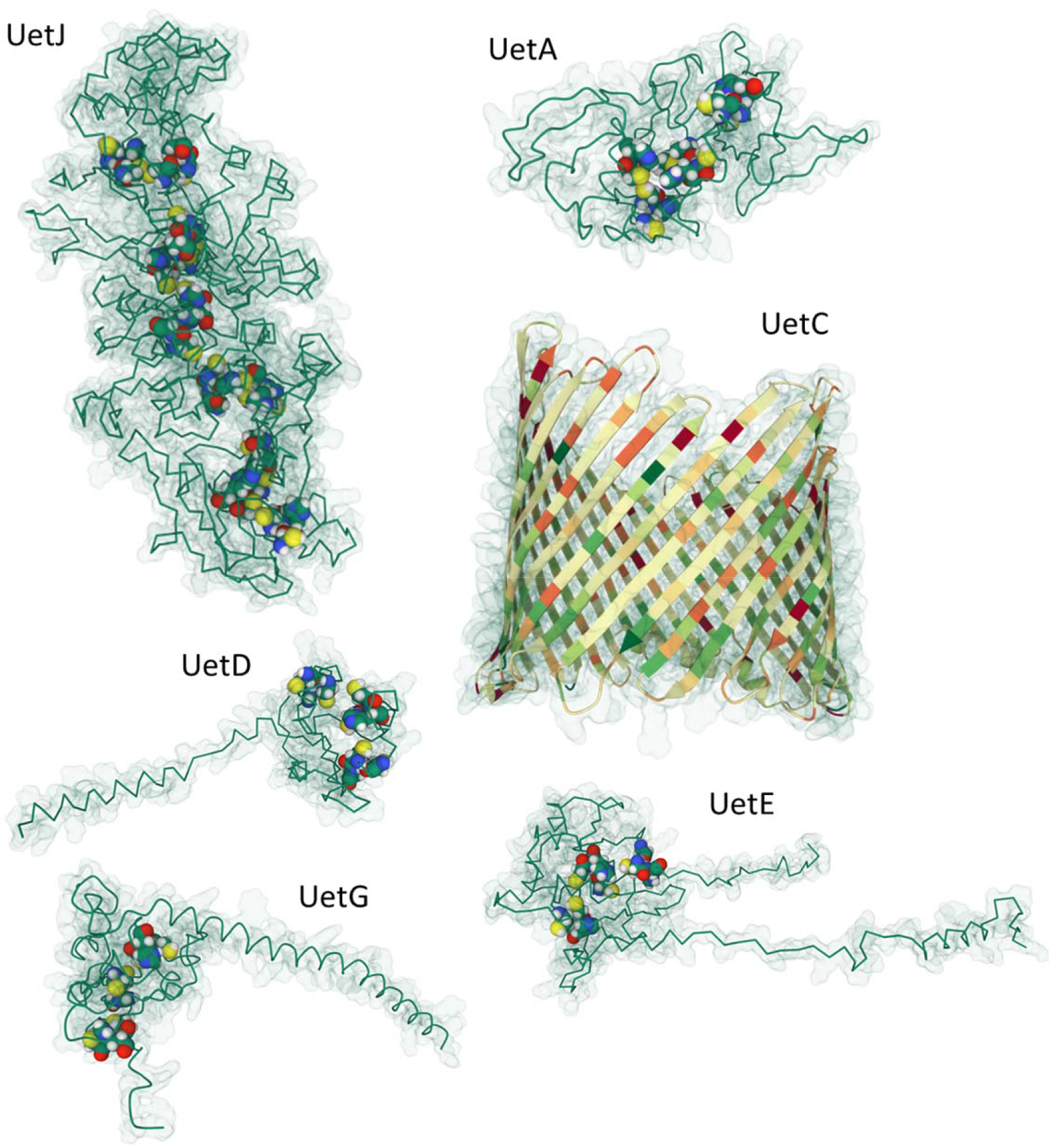
Predicted protein structures. Protein backbones are shown as pipes, with calculated Gaussian surface shown as transparent green. Heme binding motifs are shown as space-filled atoms to highlight the position of hemes. UetC is shown with a ribbon cartoon, with β-strands shown as flat arrows, and colored by hydrophobicity with red residues indicating hydrophilic and green indicating hydrophobic.

Supplemental Table 1: Full genome set statistics and extended data. Colors highlight groups as displayed in Figure 1, Supplemental figure S1, Supplemental figure S2.

Supplemental Table S2. Membership of orthogroups. Orthogroup identifiers (column D) are linked to member orthologs (columns E – EA) colored by genome group as seen in Figure 1, Supplemental figure S1, Supplemental figure S2.

Supplemental table S3. Numerical membership of all orthogroups (columns A) with number of representatives (columns B-F), and proportional prevalence (columns G-K).

Supplemental table S4. Orthogroups described in this paper and functional predictions. Automated gene annotations from NCBI or RAST are shown in Column C, predicted domains, features and functions in column D, predicted role or complex column E, localization column F, number of representatives in columns G-K, and proportional prevalence in columns L-P with colors shown in supplemental table S3.

